# Developing a Biomimetic Evaluation Method for Antiviral Coatings Using Artificial Saliva Droplets

**DOI:** 10.1101/2021.10.21.465373

**Authors:** Naoki Tanaka, Nobuhiro Miyamae

**Affiliations:** Nippon Paint Co., Ltd., 4-7-16 Minami-Shinagawa, Shinagawa, Tokyo 140-8677, Japan

**Keywords:** antiviral agents, viral infections, M13 phage, biomimetic evaluations, antiviral coatings

## Abstract

Respiratory infections pose a serious threat worldwide, and many new antiviral agents and coatings have been developed to reduce the overall risk of viral infection. Here, we evaluate the methodology used to test these antiviral coatings and developed a novel system that is more similar to “real-world” conditions. Contact infection is largely mediated via contact with saliva containing the active virus released as droplets by coughing or sneezing, with these droplets adhering to objects and surfaces and subsequently entering the human body via indirect contact with the mucous membranes. Here, we evaluated the antiviral effect of a known antiviral coating agent using an artificial saliva based system, where artificial saliva containing phages were sprayed onto the antiviral coating under various conditions associated with viral replication and infectious spread. We used a commercially available antiviral coating in this evaluation, and M13 bacteriophages as model viruses. This method enables simple biomimetic evaluations of any product’s antiviral effects.

## Introduction

Recent outbreaks of viral diseases have highlighted the risks associated with both seasonal viral infections, such as influenza and noroviruses, and epidemic-level respiratory infections such as COVID-19, caused by the SARS-CoV-2; thus, the prevention of such infections has become a top priority ^[1-3]^. Respiratory pathogens can infect the human body via numerous routes including contact, airborne infection, or droplet-based infection, with droplet-based contact infections being the most common ^[4, 5]^. Contact infections are mediated by the active viral particles contained in the saliva released from infected persons as droplets during coughing or sneezing ^[6, 7]^; these viral particles adhere to many different surfaces including handrails, doorknobs, walls and push buttons, and subsequently enter the human body via indirect contact with the mucous membranes ^[8- 11]^.

Various surface treatments containing antiviral agents have been developed to reduce the risk of viral infection via this type of community spread ^[12-14]^ and the evaluation of these coatings are generally standardized by the International Organization for Standardization ^[15, 16]^. These methods typically involve a predetermined set of temperature and humidity conditions and generally evaluate bacteriophage Qβ, influenza viruses, or feline caliciviruses. A recent study described a novel evaluation method that relies on the use of the M13 bacteriophage and phagemid systems ^[17]^.

However, these methods do not replicate the actual environment where these infections occur. For example, conditions such as the virus dispersion medium, temperature, and humidity in real infectious environments may differ significantly from the general evaluation method. These shortcomings were recently addressed for antimicrobial evaluation methods ^[18]^, but this method is not applicable to viruses. A recent evaluation for an anti-SARS-CoV-2 coating material was evaluated using a more *in vivo*-like environment and helped to determine the effects of sebum on high-touch surfaces on antiviral coatings ^[19]^, however, the effect of contaminated saliva droplets was not evaluated. Given these shortcomings, we designed this study to develop a novel antiviral evaluation method which more closely resembles real-world infections.

Our study used the commercially available PROTECTON BARRIAX™ SPRAY (Nippon Paint Industrial Coating Co., Ltd.) antiviral coating as it has been shown to have antiviral effects against influenza viruses and bacteriophage Qβ, as shown in Fig S1. Furthermore, we also focused on the safety and efficiency of our evaluation designed and we completed our evaluations using M13 bacteriophages which is a known reference model for these kinds of evaluation ^[17]^.

## Materials and Methods

### Preparation of M13 bacteriophage stock solutions

M13 phage stock was prepared as previously described ^[20]^. Briefly, the *Escherichia coli* stock containing an antibody library was grown in broth and infected with helper phages before being cultured overnight and then subjected to PEG precipitation and dissolved in SM buffer (10 mM Tris-HCl pH 7.5, 100 mM NaCl, and 8 mM MgSO_4_). Purified phages were then stored at 4 °C and the number of active phages in the solution was established to be 10^11^–10^12^ CFU/mL.

### Preparation of phage suspension for antiviral coating evaluations

We used artificial saliva as the phage dispersion medium in our novel evaluation method with this solution being prepared as previously described ^[21]^. We used 42.4 mg of sodium chloride, 60.0 mg potassium chloride, 7.3 mg calcium chloride hydrate, 2.6 mg magnesium chloride, and 17.1 mg dipotassium phosphate, as the inorganic component and 125 mg of porcine mucin (137-09162; FUJIFILM Wako Pure Chemical Corporation, Osaka, Japan) mixed in 50 mL of pure water as the organic component. The phage stock solution was diluted 10-fold using artificial saliva and used for evaluation. We also produced a 10-fold phage stock solution using phosphate-buffered saline (PBS) to allow for a clear comparison of the two evaluation methods.

### Evaluation of phage inactivation by the antiviral coating

Our inoculation method is clearly explained in Fig. 1. Briefly, we inoculated 50 µL of each M13 phage suspension onto glass slides (50 mm × 50 mm) with or without the antiviral coating and covered them with 0.06 mm thick transparent polypropylene film (40 mm × 40 mm), as shown in Fig. 1[a]^[17]^, referred to as the “previous method” in this report. Artificial saliva was dispersed onto similarly treated slides using a spray bottle held 15 cm away from the slide (Fig. 1[b]), which is described as the “spray method” in this report. After incubation under fluorescent light irradiation at 500 lx, M13 phages were collected by rinsing in 1000 volumes of PBS or artificial saliva. The eluted and serially diluted phages were then mixed with *E. coli* strain XL-1 Blue cells at OD600 range = 0.4–0.6 and incubated in a water bath at 37 °C for 30 min. Then 5 μL of each *E. coli* cell was spotted into grids on TYE (Triptone Yeast extract agar) plates containing ampicillin and incubated overnight at 37 °C before the number of colonies were counted and the number of active phages were determined.

**Fig. 1.**
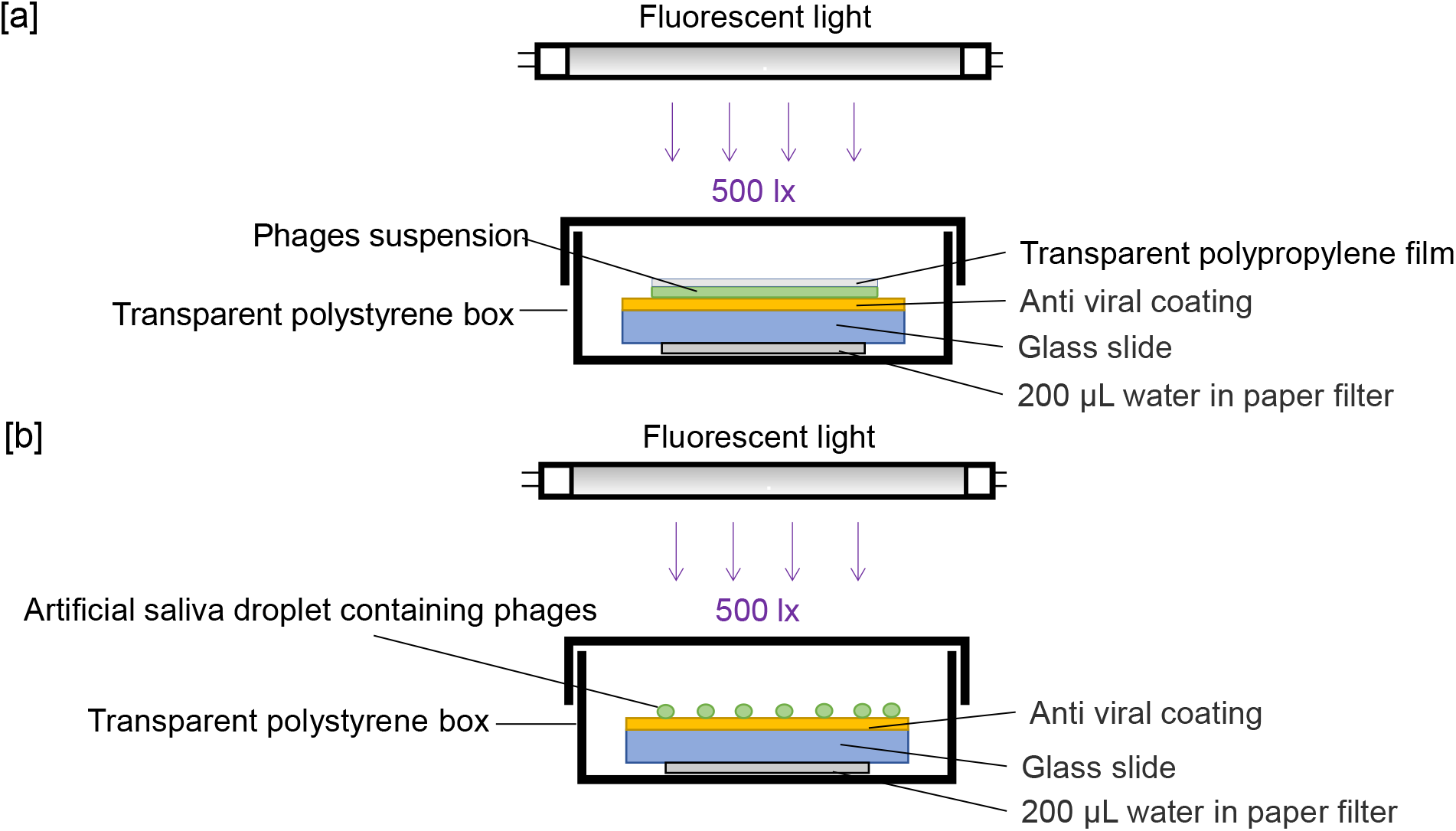
Schematic describing phage inoculation. [a] Illustrates the classical inoculation method used in previous evaluations. Here, a glass slide is placed on filter paper saturated with 200 μL of sterilized water and then inoculated with a phage suspension before being covered with a transparent polypropylene film. [b] Describes our novel method of inoculation, where a similar glass slide is placed on filter paper saturated with 200 μL of sterilized water and then treated using a spray bottle to emulate the spray based dispersal associated with saliva, creating a more biomimetic evaluation. Droplets were dispersed using a commercially available glass spray bottle.

## Results

We have tried to develop a novel antiviral evaluation method that more closely resembles the actual environment in which viruses may be spread, under various conditions associated with viral replication and infectious spread, such as saliva droplets containing viruses, time course of virus titer and temperature and humidity. Fig. 2 shows the particle size distribution of the artificial saliva droplets produced using the spray method replicating what we expect from a cough or sneeze onto a coating, revealing that they had a mean diameter of 207 μm, a median diameter of 51 μm, and a mode of 69 μm when dispersed using the spray method. Fig. 3 shows the viral titers of the M13 phages in each of the droplets produced using the spray method replicating what we expect from a cough or sneeze onto an antiviral coating. The phage titer for the control was shown to be 2.4 × 10^9^ CFU/mL and under the limit of detection for the treated glass when evaluated using the traditional droplet method, while the control slides produced a titer of 9.1 × 10^9^ CFU/mL when using the new spray method. Notably, this more biologically relevant dispersal produced a titer of 2.8 × 10^5^ CFU/mL on the antiviral coated slides, suggesting that while both methods allow for the evaluation of efficacy, the spray method is more robust and likely more representative of the real world. In addition, both were inactivated by more than 99.9% and met the significance level with p-values of 0.0002 and 0.00003, respectively. This indicated that the spray method could be used for proper antiviral evaluation. Fig. 4 illustrates the change in viral titer overtime when surfaces are exposed to more environmentally relevant conditions. Both methods revealed that producing inactivation rates of 95.2% at 10 min and 98.9% at 30 min when compared to the control, the phage titer decreased over time in response to the antiviral coating. Fig. 5 shows the change in viral titer over time in response to different temperature and humidity conditions. The higher the temperature, the faster the viral titer decreased while this was not as obvious for humidity. These results suggest that M13 phage titer was more sensitive to temperature than humidity.

**Fig. 2.**
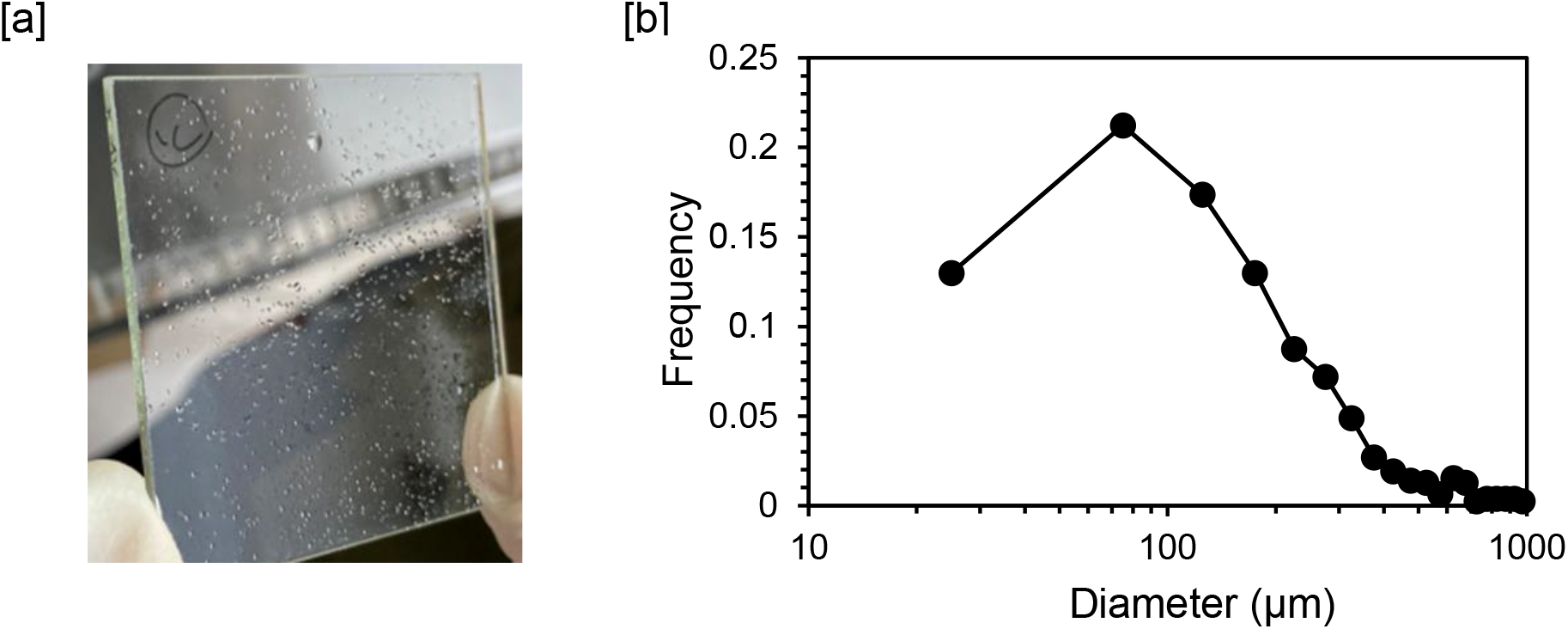
Photographic image and particle size distribution of droplets made using a commercial sprayer. The droplets on the glass slides were photographed using a digital microscope (VHX-7000, Keyence Corporation) and the particle size was measured. [a] Is a representative photograph of the artificial droplets on the glass slide and [b] is a graph describing their particle size distribution. The mean diameter was 207 μm, median diameter was 52 μm, and mode was 70 μm.

**Fig. 3.**
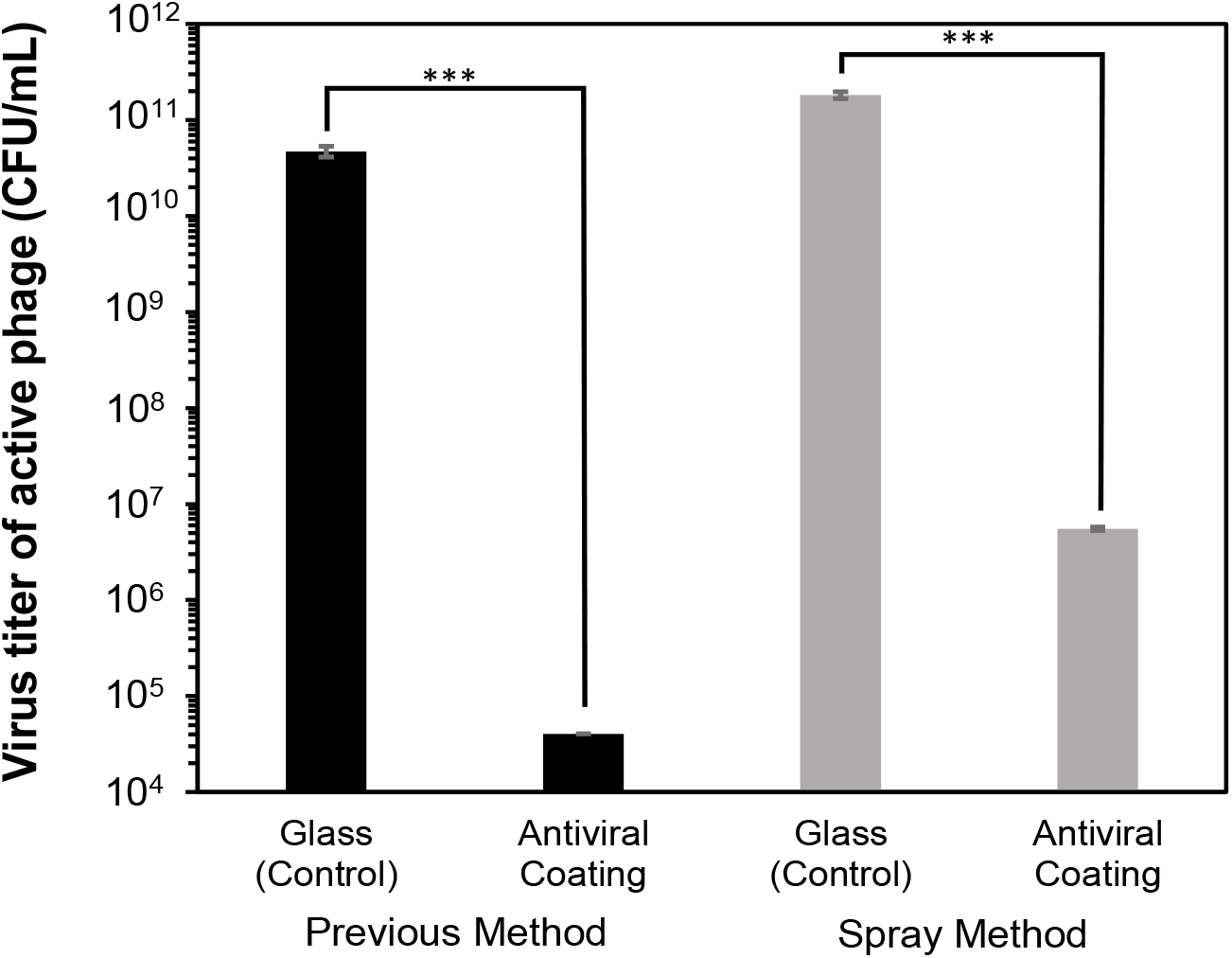
Viral titers of the M13 phages. We inoculated a blank (control) or treated glass slide (antiviral coating) with M13 phage solution and evaluated the antiviral efficacy of our novel antiviral coatings using a colony forming unit assay. The suspension used in one set of samples was covered with a polypropylene film as described in the earlier inoculation methods, while the other set of inoculated slides were treated with artificial saliva using the “spray method”. Data is represented as the mean ± S. D., n= 3 per a group. and the p-values were calculated using one-way ANOVA where * p < 0.05, ** p < 0.01 and *** p < 0.001.

**Fig. 4.**
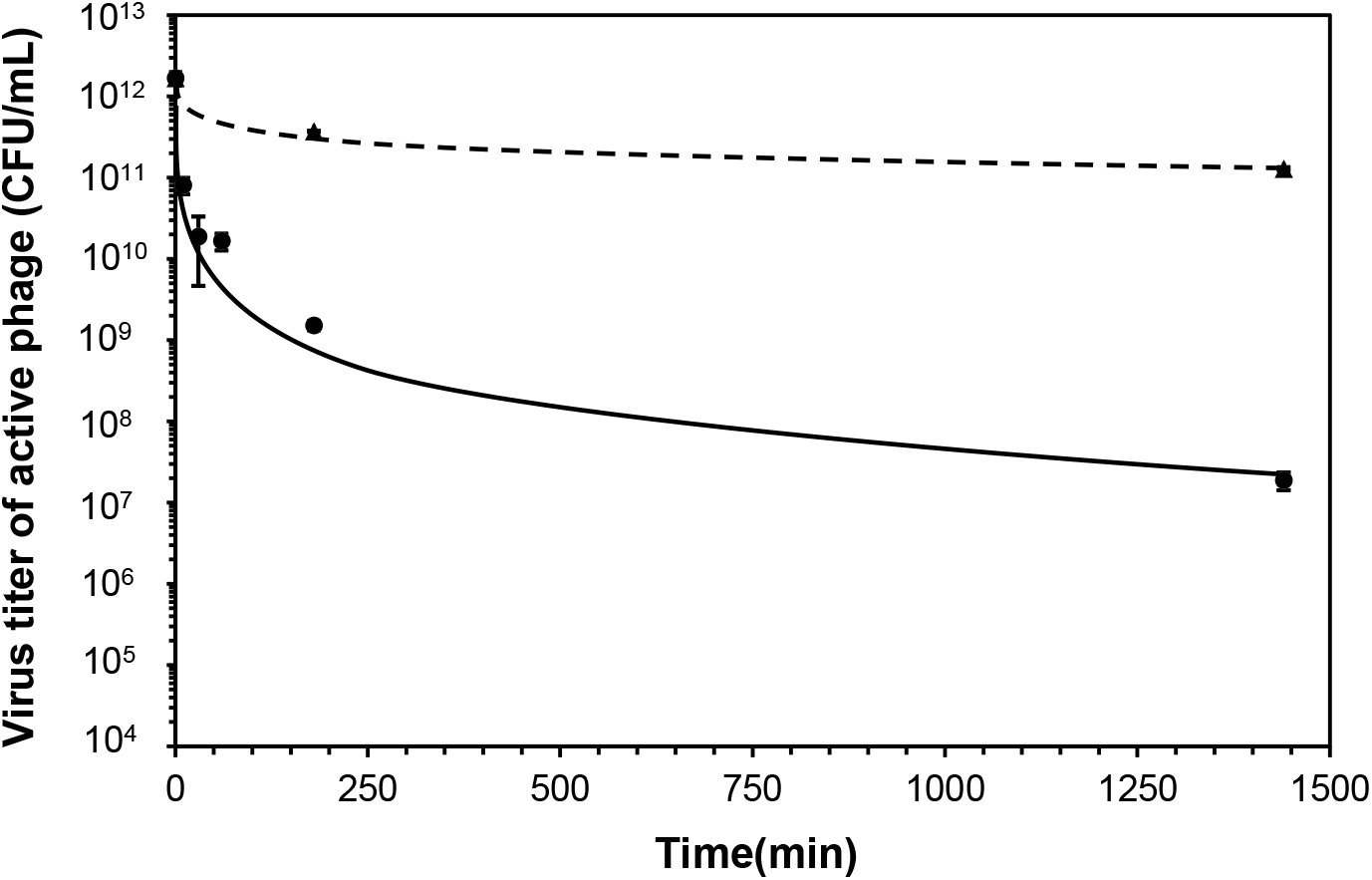
Time-course evaluating the phage titer when inoculated using the spray method. The active phage titer for various time points (10, 30, 60, 180, and 1440 min) was evaluated at 23 °C. The dotted line represents the control slides while the solid line represents the results from the coated slides. Values represent the mean ± S. D., n= 3 per group.

**Fig. 5.**
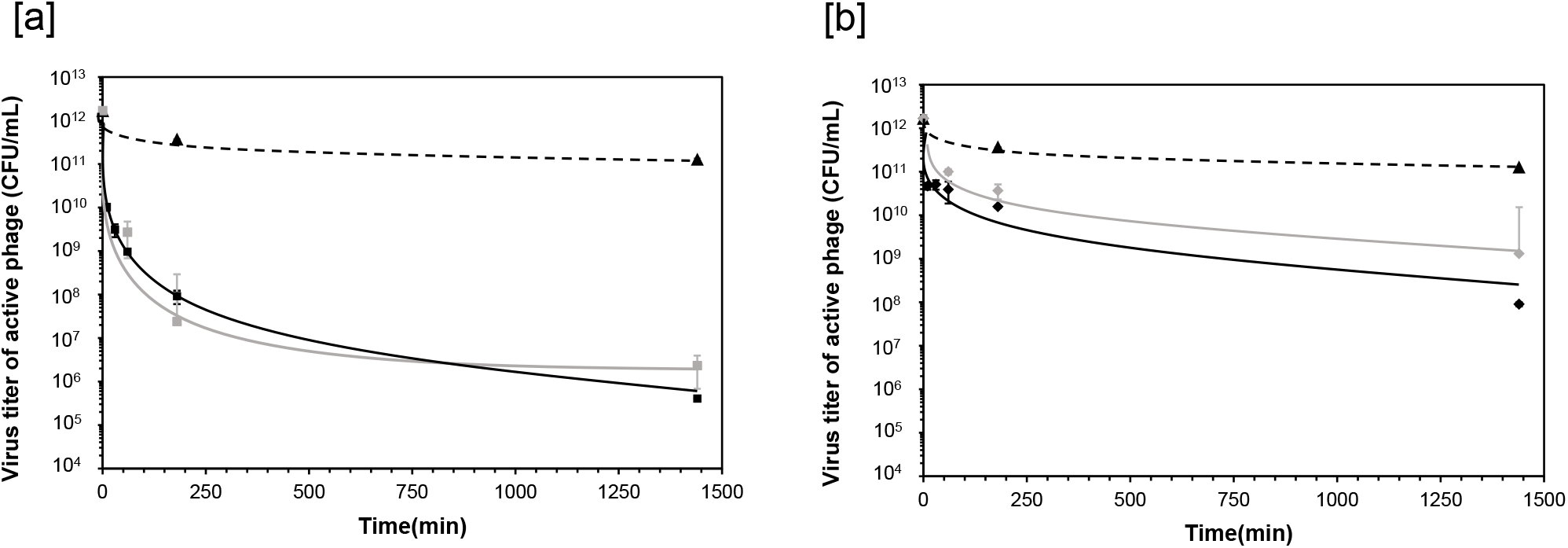
Time-course describing changes in phage titer under different temperature and humidity conditions. [a] Time-course for active phage titers at 40 °C and [b] 10 °C. The dotted lines in each graph represent the results from the control, and the solid lines represent the results for the antiviral coating. In addition, black lines indicate results at a humidity of 80% and gray lines a humidity of 25%. Temperature and humidity were controlled using a water bath and silica gel, respectively. All data are represented as the mean ± S. D., n= 3 per group.

Thus, it revealed to have slightly difference in the evaluation results between previous method and the spray method, but the spray method was shown to be effective as a method for antiviral evaluation more closely resembles the actual environment in which viruses may be spread.

## Discussion

We have tried to develop a novel antiviral evaluation method that more closely resembles the actual environment in which viruses may be spread, under various conditions associated with viral replication and infectious spread. The particle size distribution of the droplets generated by coughing, sneezing, or speech is approximately 10–2000 μm ^[22-24]^. The particle size distribution of the droplets created using the spray method in this study fall within this range, suggesting that they are more likely to reflect the environmental samples more accurately. The major differences between the more *in vivo-*like environment created in this study and the previously described test conditions center around viral dispersion and the differences between the spray and droplet mediated viral distribution on test surfaces. As Fig. 3 shows, the number of active phages were significantly reduced when brought into contact using either dispersal method, but that this decrease was less significant in the spray method, suggesting its superiority when evaluating antiviral effects. In addition, active phages were shown to be rapidly inactivated upon contact with the antiviral coating (Fig. 4), and that even more resistant phages became inactivated over time. In addition, the higher the temperature at the time of inoculation, the more the virus was inactivated when applied using the spray method, which was similar to the results reported by several previous studies ^[25, 26]^. We speculate that the higher the temperature, the more the antiviral agent on the coating will denature the proteins the virus uses to bind to its host, reducing infectivity. Thus, we suggest that the higher the temperature, the higher the antiviral activity of the coating agent in the actual environment.

The viscosity of artificial saliva, which was used as the phage dispersion medium in this study, was evaluated using a BL-type Viscometer (TOKYO KEIKI, INC.), and the effect of each dispersal method on the relative antiviral properties of the treatment was then confirmed. The viscosity of artificial saliva (25 °C) used in this study was 3.0 mPa·s. It is also generally known that the viscosity of water (25 °C) is 0.8 mPa.s ^[27]^. The reason for the high viscosity of artificial saliva is thought to be associated with the use of mucin, which is a mixture of glycoproteins, to produce artificial saliva. The flow of the M13 phages dispersed in these droplet solutions suppressed because the viscosity of the artificial saliva was higher than that of water. Therefore, the spray method using artificial saliva reduced the amount of contact between the M13 phage and the antiviral coating. It is also known that the viscosity of human saliva varies depending on a person’s physical condition, activity, and age ^[28-31]^. The artificial saliva used in this study was adjusted to reflect the average viscosity of human saliva and future work should include some evaluation of the antiviral activity of these coatings using saliva of varying viscosity.

We also measured the shape of the artificial saliva, which was sprayed on the glass and dried using a non-contact optical profiler (CCI MP-HS, AMETEK Inc.). The mucin-containing artificial saliva forms a layer on the glass during the drying process (Fig. 6), suggesting that those M13 phages incorporated into the mucin layer are prevented from interacting with the antiviral coating.

**Fig. 6.**
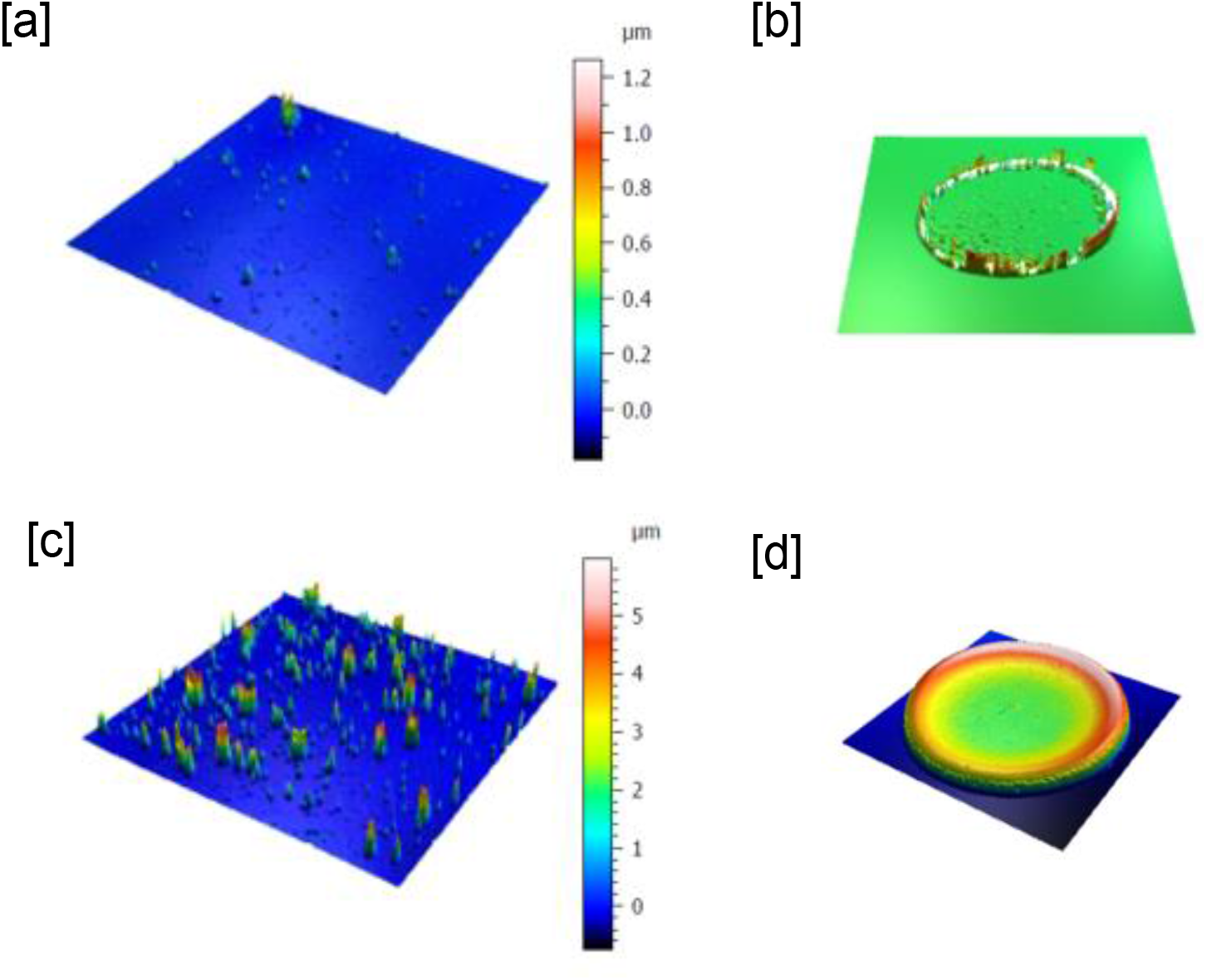
The shape of dried PBS and artificial saliva droplets. Droplet shape was evaluated using a non-contact optical profiler (CCI MP-HS, AMETEK Inc.) and [a] shows the surface roughness of PBS buffer droplets while [b] shows the shape of the dried PBS buffer. The center of the dried PBS droplets is almost 0 μm while the thickness at the edge is approximately 1 μm. [c] Shows the surface roughness of dried artificial saliva on glass whole [d] shows the shape of the dried artificial saliva droplets. These drops are approximately 2 μm thick in the center and 5 μm thick at the outer edge.

In the future, it will be necessary to visualize the amount of virus incorporated into the mucin layer. Although there are still several parameters that need to be addressed to produce more reliable *in vitro* evaluations of these coating products, the use of the spray dispersal method does seem to allow for a closer recapitulation of the natural environment and as such is expected to become a standard practice in the evaluation of these surface treatments.

## Conclusion

We have developed a novel antiviral evaluation method that more closely resembles the actual environment in which viruses may be spread, facilitating the better evaluation of these products for reducing the risk of contact infection. This report shows that the selection of the virus dispersant and dropping method of the viral suspension are both important factors in simulating the actual infectious environment. Thus, the antiviral effect can be more accurately evaluated using the spray method and artificial saliva containing the virus of interest (in this case M13 bacteriophages as a model virus). We also revealed that inoculation temperature had a significant effect on the inactivation of M13 bacteriophages, and suggest that the spray method using artificial saliva containing M13 bacteriophages is safe and simple, and is expected to facilitate more biologically relevant evaluations of antiviral coatings.

## Acknowledgements

We wish to thank Makoto Nakakido and Professor Kouhei Tsumoto from the School of Engineering at Tokyo University for their useful advice and technical assistance during our experiments. We would like to thank Editage (http://www.editage.com) for their manuscript review and editing support.

