## Supplementary Material.pdf for "Developing a Biomimetic Evaluation Method for Antiviral Coatings Using Artificial Saliva Droplets"

Supplemental Figure

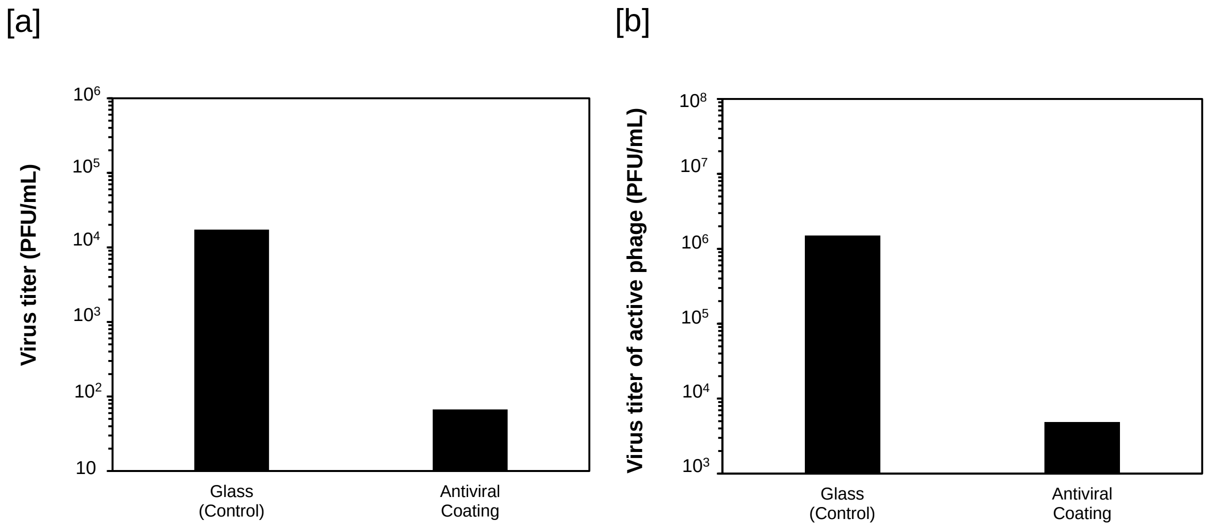


Fig. S1 Antiviral effect of commercially available antiviral coating. [a] Antiviral effect of PROTECTON BARRIERX SPRAY on Influenza. This result was first published in report No. 20077016001-0201 from the Japanese Food Research Laboratories (JFRL). [b] Shows the antiviral effect of PROTECTON BARRIERX SPRAY on Qβ phages, with this result first reported in report No. PI2011R-1 from the TOTO Ltd. Research institute.

[1] Correspondent:
Naoki Tanaka
Nobuhiro Miyamae
